# Efficient axonal trafficking of endolysosomes depends on the balanced ratio of microtubule tyrosination and detyrosination

**DOI:** 10.1101/2023.10.18.562903

**Authors:** Anja Konietzny, Leticia Peris, Yuhao Han, Yannes Popp, Bas van Bommel, Aditi Sharma, Philippe Delagrange, Nicolas Arbez, Marie-Jo Moutin, Marina Mikhaylova

## Abstract

In neurons, the microtubule (MT) cytoskeleton forms the basis for long-distance protein transport from the cell body into and out of dendrites and axon. To maintain neuronal polarity, the axon initial segment (AIS) serves as a physical barrier, separating the axon from the somatodendritic compartment and acting as a filter for axonal cargo. Selective trafficking is further instructed by axonal enrichment of MT post-translational modifications, which affect MT dynamics and the activity of motor proteins. Here, we compared two knockout mouse lines lacking the respective enzymes for MT tyrosination and detyrosination and we found that both knockouts led to a shortening of the AIS. Neurons from both lines also showed an increased immobile fraction of endolysosomes present in the axon, whereas mobile organelles displayed shortened run distances in the retrograde direction. Overall, our results highlight the importance of maintaining the balance of tyrosinated/detyrosinated MT for proper AIS length and axonal transport processes.

**Summary Statement:** Despite opposite effects on microtubule dynamics, shifting the balance of microtubule tyrosination/detyrosination in either direction resulted in surprisingly similar defects in axonal organelle transport.

## Introduction

The complex, polarized morphology of neurons with their extensively branched neurites presents various challenges for the maintenance of cellular homeostasis. Neurons contain an elaborate intracellular recycling system within the soma, dendrites and axon that balances the transport of newly synthesized cellular components with the removal of aged and damaged ones. At the same time, transport processes need to be highly selective to maintain the molecular identities of the somatodendritic and the axonal compartments. The master regulator of polarized trafficking is a complex structure spanning the first 20-60 µm of the axon, called the axon initial segment (AIS), which selectively restricts cargo transport and is essential for the generation of action potentials (Nakada et al., 2003; Petersen et al., 2014). The AIS is composed of various elements, including transmembrane receptors, ion channels, a specialized membrane-associated periodic skeleton (MPS) rich in F-actin and spectrin, parallel microtubules (MT) that are bundled and interlinked through tripartite-motif-containing 46 (TRIM46; Van Beuningen et al., 2015), as well as dynamic EB3-positive MTs. All these components are assembled by the master scaffolding protein Ankyrin-G (AnkG; Leterrier et al., 2015). Throughout the entire cell, MT tracks provide the basis for long-distance transport, including that of recycling organelles. The clearance of damaged cellular components is mediated to a large part by acidic vesicular organelles, such as late endosomes and lysosomes, as well as a range of autophagic vesicles which are present along the axon. Over the past years, disturbances in clearance pathways have been identified as a common denominator in many neurodegenerative diseases (Menzies et al., 2017; Pitcairn et al., 2019; Root et al., 2021; Wang et al., 2018), with axonal damage emerging as a major driver of neurodegeneration. In healthy axons, endolysosomes delivered from the cell body ensure the timely clearing of damaged proteins (Farfel-Becker et al., 2019), and participate in signaling and RNA trafficking (Liao et al., 2022; Napolitano et al., 2019). Such MT-based transport into the axon is driven by motor proteins whose activity is regulated by selective MT post-translational modifications (PTMs)(Park & Roll-Mecak, 2018). Abundant MT PTMs in neurons are tubulin tyrosination and detyrosination. Tyrosinated MT are generated from *de novo* synthesized α-tubulin, which carries the C-terminal tyrosine residue, or by tyrosine ligation to detyrosinated tubulin by the tubulin-tyrosine ligase (TTL; Ersfeld et al., 1993). MT tyrosination has been identified as a marker for dynamic microtubules (Moutin et al., 2021; Tas et al., 2017). MT-detyrosination, which is largely catalyzed by the tubulin-detyrosinase complex vasohibin1/2-small vasohibin binding protein (VASH-SVBP), occurs predominantly in the axon, where it marks a long-lived and stable MT pool (Fig. 1A; C. et al., 2017; Moutin et al., 2021). Plus-end directed kinesin motors (Cai et al., 2009; Dunn et al., 2008; Gumy et al., 2013; Homma et al., 2000; Konishi & Setou, 2009; Peris et al., 2009), as well as minus-end directed dynein motor complexes (McKenney et al., 2016; Nirschl et al., 2016; Peris et al., 2006) are known to be sensitive to the tyrosination/detyrosination (Tyr/deTyr)-state of MTs, suggesting a role of MT tyrosination in the regulation of axonal molecular transport.

**Figure 1:**
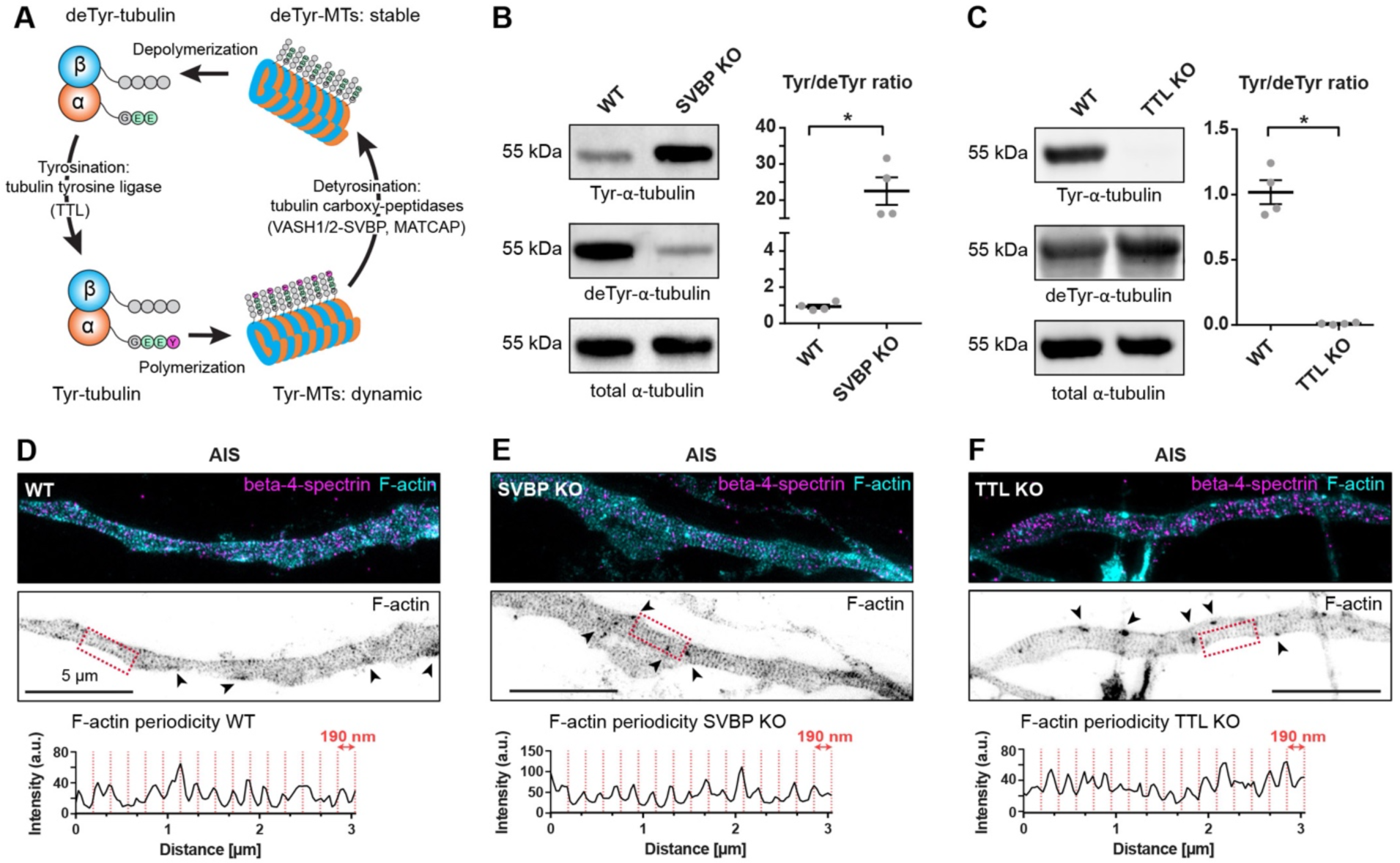
Loss of MT tyrosinating/detyrosinating enzymes does not affect the nanostructure of the F-actin cytoskeleton in the AIS. **A)** Illustration of the tubulin tyrosination/detyrosination cycle. **B, C**) Immunoblot quantifications of the ratio of tyrosinated vs. detyrosinated tubulin from crude lysates of cultured mouse cortical neurons at DIV15 with SVBP-KO (B) and TTL-KO (C) compared to WT neurons. n = 4, 3 independent cultures for WT, SVBP-KO respectively, and 4, 3 independent cultures for WT, TTL-KO respectively. Kruskall-Wallis test with Dunn’s multiple comparison test, ns = non-significant, *p<0.05. **D-F**) STED images of the AIS of DIV12 neurons stained with phalloidin-647 (F-actin marker) and with an antibody against beta-4-spectrin. Arrows indicate F-actin patches. **D)** In wildtype neurons, F-actin and beta-4-spectrin form an intercalated submembrane scaffold complex with a 190 nm periodicity. **E, F)** MPS is formed normally in both TTL (B) and SVBP-KO (C) neurons.

Here, we used two previously established knockout (KO) mouse models, TTL-KO and SVBP-KO, to characterize the influence of the microtubule Tyr/deTyr status on the structure of the AIS as well as on the axonal transport of acidic endolysosomal organelles. We found that TTL- and SVBP-KO both resulted in an overall shortening of the AIS, without affecting the nanostructure of the MPS. Furthermore, axons of both TTL- and SVBP-KO neurons contained a higher fraction of immobile endolysosomes, with the mobile fraction displaying markedly decreased run lengths in the retrograde direction. Thus, although VASH-SVBP and TTL activity are known to have opposite effects on MT dynamics (Sanyal et al., 2023), we found that the loss of either enzyme affects AIS structure and axonal trafficking in a similar way, indicating that the regulation of axonal cargo transport relies on maintaining a fine balance between MT tyrosination and detyrosination.

## Results and discussion

### Alterations of the microtubule Tyr/deTyr status are associated with shortened AIS without affecting F-actin organization

We first quantified the tubulin Tyr/deTyr ratios via immunoblot analysis of tyrosinated, detyrosinated and total α-tubulin in primary wildtype (WT), SVBP-KO and TTL-KO cultured cortical neurons. As expected, this ratio was increased in SVBP-KO neurons (22.540±3.804) and dramatically reduced in TTL-KO neurons (0.010±0.003) (Fig. 1B,C). It was previously shown that neurons of TTL-KO mice exhibit premature axon differentiation and formation of supernumerary axons, suggesting that enrichment of stable, detyrosinated MTs induces axon formation (Erck et al., 2005; Peris et al., 2009). In contrast, SVBP-KO neurons, devoid of VASH-SVBP carboxypeptidases, exhibit a delay in their axon development and reduced axonal length (Aillaud et al., 2017; Pagnamenta et al., 2019). To investigate whether this would also have deleterious effects on the AIS structure, we first conducted stimulated emission-depletion (STED) imaging in primary hippocampal neurons to visualize the F-actin-spectrin MPS (Leterrier et al., 2015). We found that neither an increase (SVBP-KO) nor a reduction (TTL-KO) of the Tyr/deTyr ratio interfered with the formation of the MPS or the presence of F-actin patches in the AIS (Fig. 1D-F). Therefore, it is unlikely that the barrier function of the AIS, which relies on the F-actin cytoskeleton (Song et al., 2009; Watanabe et al., 2012), is affected. However, polarized trafficking is additionally instructed by uniformly oriented axonal MTs, which are bundled tightly by the protein TRIM46 inside the AIS (Van Beuningen et al., 2015). We therefore decided to compare the distribution of two AIS markers, AnkG and TRIM46, in proximal axons of KO and WT mouse hippocampal neurons using a MATLAB script based on previously established criteria (Grubb & Burrone, 2010; Fig. 2A). Representative images of WT and KO neurons stained for TRIM46 (and AnkG for WT) are presented in Fig. 2 B-E. In TTL-KO neurons, both markers indicated significantly shortened AIS (AnkG: 22.1±0.3 µm, TRIM46: 19.3±0.5 µm) compared to WT (AnkG: 29.1±0.8 µm, TRIM46:

**Figure 2:**
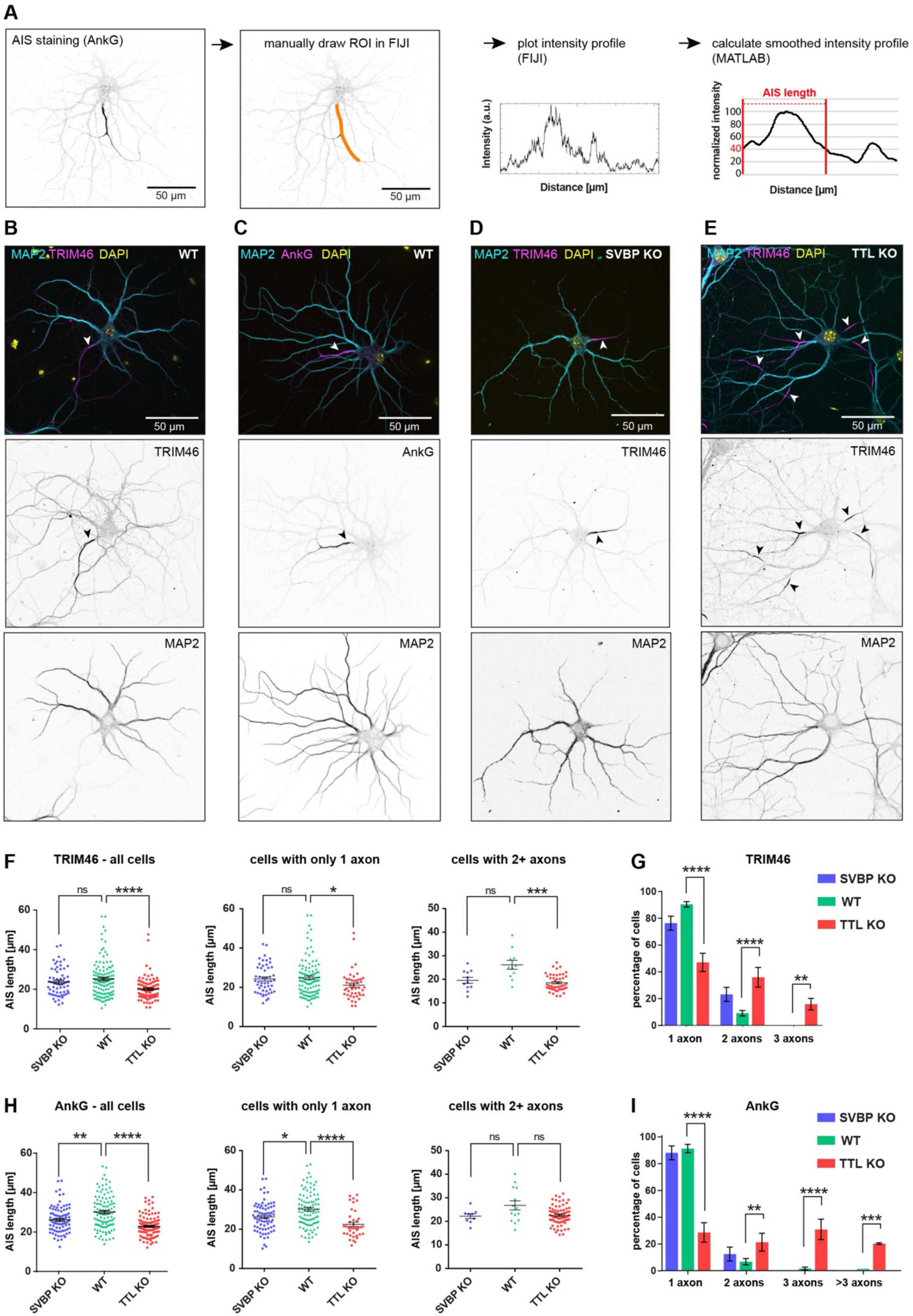
Both SVBP and TTL-KO lead to shorter AIS in hippocampal neurons. **A)** Overview of the workflow for AIS length calculation (see Material and Methods). **B, C)** Example images of WT DIV12 mouse hippocampal neurons stained for the AIS markers TRIM46 (B) or AnkG (C) together with MAP2 (dendritic marker), and DNA (DAPI). Arrows indicate the AIS. **D, E)** Representative images of DIV12 TTL-KO (D) and SVBP-KO (E) neurons stained for MAP2, TRIM46 and DAPI. Arrows indicate AIS. **F, H)** AIS length of TRIM46-stained (F) and AnkG-stained (H) neurons from SVBP-KO, WT, and TTL-KO mice. **Left panel:** graphs show all cells analyzed together. **Middle, Right panel:** Graphs show cells with only 1 axon and cells with 2+ axons analyzed separately. **G, I)** Quantification of neurons with supernumerary axons in the different genotypes as judged by TRIM46 (G) and AnkG (I) staining. TRIM46: n = 68, 93, 123 AIS for SVBP-KO (2 independent cultures), TTL-KO (5 independent cultures) and WT (6 independent cultures). AnkG: n = 84, 110, 108 AIS for SVBP-KO (3 independent cultures), TTL-KO (4 independent cultures) and WT (5 independent cultures). F,G) Kruskal-Wallis test with Dunn’s multiple comparisons test. H,I) Two-way repeated measures ANOVA with Dunnett’s multiple comparisons test. J,K,L) Mann-Whitney-Test. ns = non-significant, *p<0.05, **p<0.005, ***p<0.0005, ****p<0.0001.

25.2±0.8µm; Fig. 2F&H, left). The SVBP-KO showed a similar effect on the AnkG (25.5±0.7 µm; Fig. 2H, left), and a non-significant trend towards shorter AIS with TRIM46 (22.9±0.8 µm; Fig. 2F, left). In line with previous observations (Erck et al., 2005), the absence of TTL resulted in a strong supernumerary axon phenotype – more than 50% of the analyzed neurons had two or more AIS emerging from soma or dendrites, while the SVBP-KO produced no such effect (Fig. 2E, G, I). The presence of supernumerary axons was not a determining factor in the reduction of average AIS length, since it occurred both in cells with only one axon, and cells with supernumerary axons (Fig. 2F, H, middle and right).

Interestingly, changes in the MT Tyr/deTyr state affected the distributions of TRIM46 and AnkG differently. TRIM46 seemed to be more sensitive to an increase in deTyr/Tyr MT ratios (TTL-KO), whereas AnkG was strongly affected by both increased and decreased deTyr/Tyr (TTL-KO and SVBP-KO, respectively). Both TRIM46 and AnkG are essential for the formation of the AIS during neuronal development. TRIM46 is a very early marker for neuronal polarization and localizes to the future axon even before axon specification and before AnkG clustering (Van Beuningen et al., 2015). TRIM46 is known to be carried into the AIS by the kinesin-2 motor complex (Ichinose et al., 2019), which is sensitive to the MT Tyr/deTyr state (Gumy et al., 2013), so the observed alteration of TRIM46 recruitment to the AIS could be mediated by a less efficient transport in the TTL-KO condition. However, the factors that delimit the length of the TRIM46-associated bundles inside the AIS, or the extent of the AIS itself, are currently unknown (Yang et al., 2019). TRIM46 itself may favor specific MT PTMs or tubulin isotypes that are only present at the proximal axon, even though none of the modifications known to date specifically mark the position of the AIS (Janke & Kneussel, 2010).

### Microtubule tyrosination status affects axonal retrograde trafficking of endolysosomes

Despite the differences in AIS length, which were within the physiological range, we did not observe any drastic changes of AIS structure in SVBP- or TTL-KO that would indicate an impaired cargo-filtering function. To further investigate the role of the Tyr/deTyr state of axonal MTs in the trafficking of endolysosomal organelles, we used two-compartment microfluidic chambers (MFCs; Fig. 3A) and Lysotracker, a pH-sensitive dye that non-discriminately labels acidic organelles, including late endosomes, lysosomes and autophagic vesicles. As there is some debate in the literature regarding bona fide degradation-competent lysosomes being present in distal axonal compartments (Farfel-Becker et al., 2019; Lie et al., 2021), we measured the correlation between Lysotracker fluorescence and fluorescence of Magic Red - a specific substrate of the lysosomal, proteolytically active form of cathepsinB (Ni et al., 2022) which fluoresces upon proteolytic cleavage (Fig. 3C). We found that the fluorescence intensity of Lysotracker correlated strongly with the fluorescence intensity of Magic Red (R^2^ = 0.9258; Fig. 3C, D), indicating that most Lysotracker-positive vesicles in the axon were likely to contain active cathepsinB, in other words were degradatively active.

**Figure 3:**
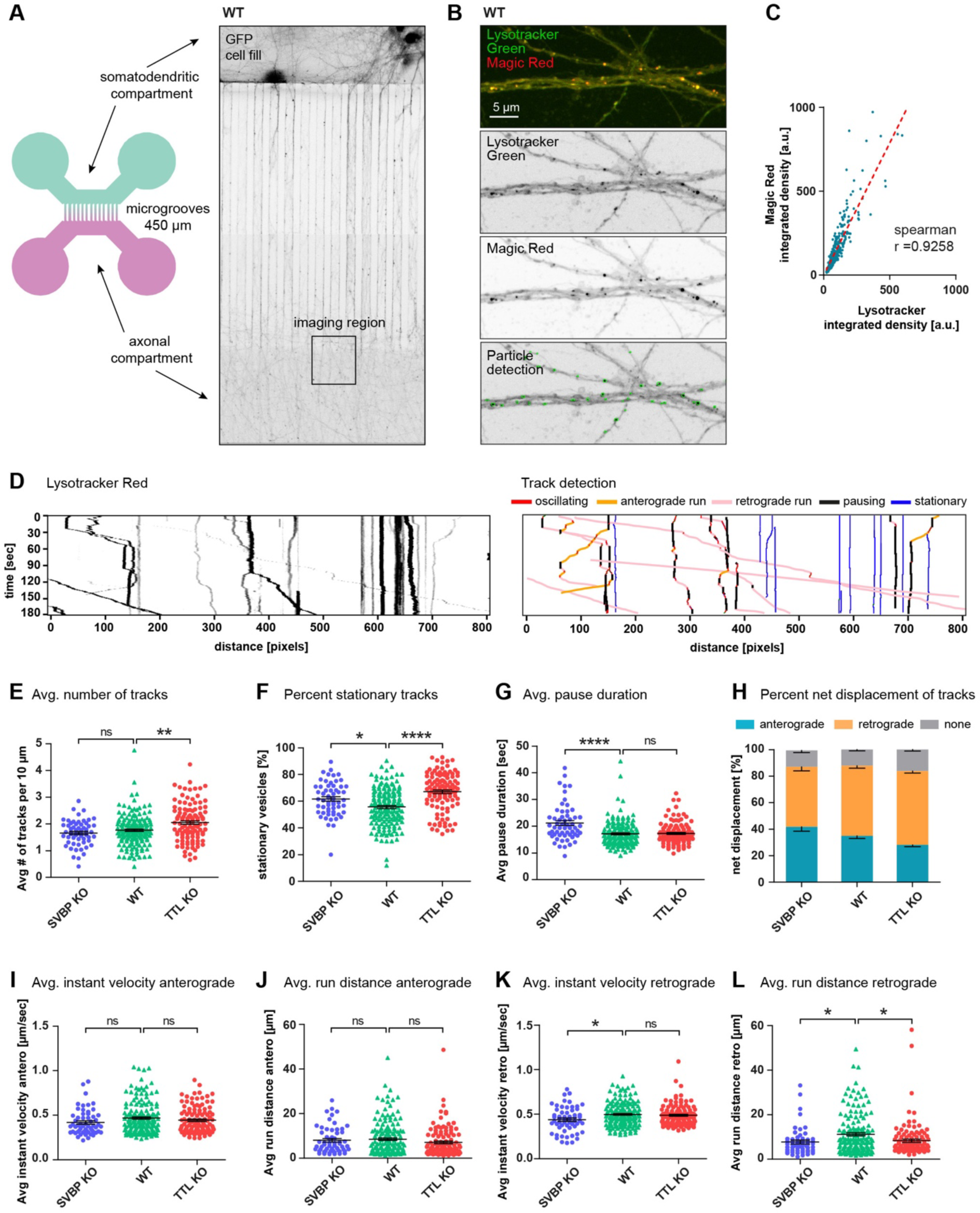
TTL- and SVBP-KO similarly reduce the efficiency of retrograde trafficking of endolysosomes in distal axons of hippocampal neurons. **A)** Illustration of the 2-chamber MFC used for live-imaging of axonal trafficking (left panel), and microscopy image of DIV12 mouse hippocampal neurons expressing GFP (right panel). **B)** Confocal image of axons inside MFC, labeled with Lysotracker Green and Magic Red. **C)** Integrated density values of Lysotracker correlate with the integrated density of Magic Red. 815 particles were analyzed in three separate locations in an area of approx. 0.2 mm². The dotted red line indicates the calculated linear regression (y = 1,578*x + 4,156), Spearman’s correlation coefficient r = 0.9258.. **D) Left panel**: representative kymograph from individual Lysotracker-treated axons; each trace represents a single organelle. **Right panel:** Kymographs analysis presented on the left using a self-made Python script (for details see Material & Methods). **(E-L)** Summary of vesicle trafficking properties during the imaging period. **E)** Average number of all detected vesicles per 10 µm (mobile and stationary). **F)** Percentage of stationary vesicles. **G)** Average pause duration. **H)** Net displacement in anterograde or retrograde direction. **I, J)** Average anterograde instant velocities and run distances. **K, L)** Average retrograde instant velocities and run distances. n = 92, 40, 55 axons from WT (6 independent cultures), SVBP-KO (3 independent cultures) and TTL-KO (5 independent cultures). B-D, F-I) Kruskal-Wallis test with Dunn’s multiple comparisons test. ns = non-significant, *p<0.05, **p<0.005, ****p<0.0001. E) Two-way repeated measures ANOVA with Dunnett’s multiple comparisons test.

We next investigated whether the Tyr/deTyr status of MTs had a differential impact on endolysosomal organelle transport by comparing the trafficking of Lysotracker-positive organelles in the axonal compartment of WT, SVBP and TTL-KO neurons (Fig. 3D, left; Movie S1). To minimize investigator bias and inconsistencies of manual analysis, we developed a semi-automated analysis workflow to analyze kymographs from individual axons (Fig. 3D, right; Fig. S2, https://github.com/HU-Berlin-Optobiology/AIS-project.git). We quantified the number of mobile and immobile vesicles, pausing times, direction of transport, velocity, and run lengths (Fig. 3E-L). In TTL-KO neurons, a significantly higher number of Lysotracker-positive vesicles was detected compared to WT (2.1±0.06 and 1.7±0.06, respectively; Fig. 3E). We observed a strong increase in the proportion of completely stationary vesicles in both SVBP (62±2%) and TTL-KO (67±1%) compared to WT (56±1%; Fig. 3F). Vesicles in SVBP-KO neurons (21.0±1.0 sec) took longer pauses than WT (17.0 ±0.5 sec), which was not the case for vesicles in TTL-KO (17.5±0.5 sec; Fig. 3G). In the axon, anterograde transport is carried out by plus-end directed kinesin motors, while retrograde movement is mediated by minus-end directed dynein motors. We analyzed the percentage of vesicles undergoing net displacement in the anterograde vs. retrograde direction and observed that a larger proportion of Lysotracker-positive vesicles moved in the retrograde (53±2% in WT) than in the anterograde direction (35±2% in WT), as previously reported by Lie et al. (2021) (Fig. 3H). No significant differences were observed between the different genotypes. Anterograde transport, including velocity (WT 0.47±0.01 µm/sec; Fig. 3I) and run distances (WT 8.5±0.6 µm; Fig. 3J), was unaffected by lack of either TTL or SVBP. Retrograde transport showed reduced velocities in SVBP-KO (0.43±0.02 µm/sec) compared to WT (0.50±0.01 µm/sec; Fig. 3K), and significantly shorter run lengths in both genotypes (SVBP-KO 7.7±0.8 µm; TTL-KO 8.3±0.7 µm, WT 11.2±0.7 µm; Fig. 3L).

Altogether our data reveal a sensitivity of retrograde trafficking of endolysosomes, mediated by the dynein/dynactin complex (Cason et al., 2021; Gowrishankar et al., 2021), towards the Tyr/deTyr state of MT. It is known that the dynactin subunit p150Glued and the dynein regulator CLIP-170 preferentially bind to Tyr-MT, which in WT neurons are enriched in the distal tip of the axon, to initiate retrograde transport (McKenney et al., 2016; Nirschl et al., 2016; Peris et al., 2006). Although an *in vitro* study has shown that the subsequent continuation of processive motility is unaffected by the Tyr/deTyr state of the MT track (McKenney et al., 2016), the interaction of p150Glued with the MT lattice promotes processive motility of the dynein-complex over a state of passive diffusion (Feng et al., 2020). Since p150Glued preferentially interacts with Tyr-MT, this could potentially explain the effects we see in TTL-KO. To our knowledge, no studies have quantified the processivity, binding affinity or velocity of dynein/dynactin on Tyr/deTyr MT. The mechanistic explanation of why we observed reduced movement speed and run length in the SVBP-KO is intriguing and remains open at this point, although it is likely to involve an altered environment of microtubule-associated proteins (MAPs; Monroy et al., 2020).

### shRNA knockdown of TTL leads to an increase in stationary Lysotracker-positive vesicles

Since defects in early neuronal polarization observed in TTL-KO neurons could potentially impact axonal trafficking at later stages, we wanted to test whether an increase in the relative amount of deTyr-MT after the establishment of neuron polarity would lead to a similar trafficking phenotype. We therefore conducted trafficking experiments where we knocked down the TTL enzyme after DIV5 in WT mouse hippocampal neurons, i.e. after axon and dendrite specification (Dotti et al., 1988). Expression of the shRNA knockdown construct reduced TTL levels in DIV12/13 neurons by 20% (Fig. S1A) and led to a significant decrease of the Tyr/deTyr tubulin ratio (to 85% of control) in transfected cells (Fig. S1B). Using this approach on neurons grown in MFCs, we conducted the same Lysotracker-trafficking experiments as described above, summarized in Fig. S1C-J. Most of the investigated parameters were similarly unaffected by TTL knockdown as in TTL-KO (Fig. S1E-I). Contrary to TTL-KO, TTL knockdown did not affect the average numbers of Lysotracker-labelled vesicles (Fig. S1C) or retrograde run distances (Fig. S1J). This could either be because the increase of deTyr MT by TTL knockdown was not severe enough to induce the same trafficking phenotype as seen in TTL-KO, or that the trafficking phenotype is indeed a result of early developmental defects in the knockout. However, following TTL knockdown, the relative amounts of stationary vesicles were significantly increased (TTL knockdown 46.7±2.8%, control 37.7±1.8%; Fig. S1D), similarly to TTL-KO (67±1% vs 56±1% in WT; Fig. 3F). This indicates that even a moderate tip in the Tyr/deTyr MT balance caused by TTL knockdown perturbed the efficient trafficking of axonal endolysosomes.

In summary, although VASH-SVBP and TTL catalyze opposite reactions, disturbing the balance of Tyr/deTyr MT in either direction had surprisingly similar effects on the investigated parameters, specifically run length in the retrograde direction and the amount of immobile organelles. The observed defects in axonal transport are most likely involved in the altered axon development described for TTL- and SVBP-deficient neurons (Pagnamenta et al., 2019; Peris et al., 2009), and hence in the neural tract abnormalities of TTL- and SVBP-KO mice (Erck et al., 2005; Pagnamenta et al., 2019). Interestingly, it was recently found that TTL expression is reduced in cases of both sporadic and familial Alzheimer’s disease, and neurons harboring the familial APP-V717I Alzheimer mutation exhibited decreased MT dynamics (Peris et al., 2022), suggesting a potentially altered regulation of MT-based cargo transport in these diseased neurons. Since the maturation of both endolysosomes and autophagosomes is tightly linked with their retrograde transport through the axon (Cason et al., 2021; Lie et al., 2021), in the future it would be interesting to investigate whether an altered Tyr/deTyr MT state affects this process and its possible contribution in neurodegenerative disease (Mohan et al., 2019).

## Material and Methods

### Animals

All experiments involving animals were carried out in accordance with the European Communities Council Directive (2010/63/EU). Experiments using tissue from wildtype C567BL/6J mice were carried out in accordance with the national Animal Welfare Act of the Federal Republic of Germany (Tierschutzgesetz der Bundesrepublik Deutschland, TierSchG) approved by the local authorities of the city-state Hamburg (Behörde für Gesundheit und Verbraucherschutz, Fachbereich Veterinärwesen) and the animal care committee of the University Medical Center Hamburg-Eppendorf, as well as of the Office for Health and Social Welfare (Landesamt für Gesundheit und Soziales, LAGeSo, Berlin, Germany) and the control of the animal welfare officers of the Humboldt University Berlin (reference number T HU-05/22). All experiments involving knockout mice were conducted in accordance with the policy of the Grenoble Institut Neurosciences (GIN) and in compliance with the French legislation. Mice homozygous for an inactivated tubulin tyrosine ligase allele (referred to as TTL-KO) were obtained as previously described (Erck et al., 2005). Mice homozygous for an inactivated small vasohibin binding protein allele (referred to as SVBP-KO) were obtained as previously described (Pagnamenta et al., 2019).

### Preparation of microfluidic chambers (MFC)

PDMS and curing agent (SYLGARD 184 Silicone Elastomer) were mixed thoroughly in a 10:1 ratio in a falcon tube, and centrifuged for 5 min at 4000 g to remove air bubbles. The PDMS was then filled into the epoxy molds and placed under vacuum for 30 min, then polymerized for 3 – 4 hours at 60°C. The cured PDMS patterns were carefully removed from the molds, the wells were cut out using a 4 mm biopsy punch, and the shape of the chambers was cut using a scalpel knife. They were then placed pattern-side down on sticky tape to protect from dust, and stored for up to two weeks. To prepare for use, the MFCs were removed from the sticky tape and vortexed in 96% ethanol for 3-5 min, then washed 3 x with MilliQ water and placed under a sterile hood for air drying. For surface activation, the MFCs were placed in a plasma cleaner, pattern-side up, together with 28 mm glass coverslips placed into 35 mm tissue-culture dishes (without lids), and treated with plasma for 30 sec. The chambers were then placed upside down on the glass coverslips as quickly as possible, then the lids were placed on the dishes and the chambers were baked for another 15-30 min at 60°C. For sterilization, the MFC inside the culture dishes were placed in a sterile hood under UV light for 20 min. They were then placed inside a 15 cm petri dish together with a piece of sterilized Whatman filter paper wetted with MilliQ water to prevent drying out of the chambers. For neuronal culture, the chambers were coated with poly-L-lysine as described below.

### Coating of Culture Dishes for Hippocampal Neurons

Glass coverslips and microfluidic chambers were coated with 1 mg/ml poly-L-lysine (Sigma #P-2636) in borate buffer (3.1 mg/ml boric acid, 4.75mg/ml borax, pH = 8.50) overnight at room temperature. The dishes were then subjected to a short rinse, a long rinse (1 hour), and 2 short rinses with sterile water. Finally, DMEM culture medium + 10% horse serum was added to the dishes and brought to 37°C. For STED imaging, high-precision coverslips (Marienfeld, 117580) were used.

### Preparation of knockout mouse hippocampal cultures

E18.5 mouse embryos were collected in sterile filtered 1x PBS. The brains were dissected, and the hippocampi were individually collected in 1x Hank’s Balanced Salt Solution (HBSS, Gibco #14185-045). Using a sterile plastic pipette, the 2 hippocampi of one embryo were taken in 2 ml of 1x HBSS. Subsequently, 200 µl of 10x trypsin (Gibco #14185-045) was added, gently agitated and incubated at 37 °C for 15 minutes without mixing during incubation. After incubation, the hippocampi were rinsed once with 1x HBSS at room temperature. The supernatant was removed, and 500 µl of DMEM (Life Technologies #31966047) + 10% horse serum (Life Technologies #26050088) at room temperature was added. Mechanical dissociation was performed by pipetting up and down using a P1000 pipette with a 200 µl pipette tip on top of the 1000 µl tip, up to a maximum of 10 times. For immunostainings, 20 µl of the 500 µl cell suspension were plated on glass coverslips in a 24-well plate containing 1 ml of DMEM + 10% Horse serum. For microfluidic chambers, 35 µl of the 500 µl cell suspension were centrifuged at 1000 g for 2 min, 30 µl of the supernatant were removed, and the cells were resuspended in the remaining 5 µl, and injected into the microfluidic chamber. The four wells of the chamber were then filled up with DMEM + 10% Horse serum. After plating, the cultures were incubated at 37°C with 5% CO2 for 2 hours, before the culture medium was replaced with MACS Neuro Medium (Miltenyi Biotec #130-093-570) supplemented with B27 (Life Technologies #17504044).

### Genotyping

PCR amplifications were performed on alkaline lysates of toe clips or tail cuts of E18.5 mouse embryos. Briefly, mouse tissue was incubated for 30 min at 95°C in alkaline solution (NaOH 25 mM, EDTA 0.2 mM, pH 12.0). Neutralization was performed by adding Tris 40 mM, pH 5.0. Lysates were then analyzed by PCR with corresponding primers and Econo Taq PLUS Green Mix (Euromedex). Primers pairs for testing TTL mouse strain were 5’GGCGACTCCATGGAGTGGTGG and 5’CCCAACATCACATTCTCCAAATATCAAAG (TTL wildtype, 1032 bp) and 5’GATTCCCACTTTGTGGTTCTAAGTACTG and 5’ CCCAACATCACATTCTCCAAATATCAAAG (TTL knockout, 900 bp). Primers pairs for testing SVBP mouse strain were 5’GATCCACCTGCCCGGAAA and 5’TTTCTTCCAGCACCCTCTCC (SVBP wildtype, 170 bp) and 5’TTTCTTCCAGCACCCTCTCC and 5’CAAACCATGGATCCACGAAA (SVBP knockout, 167 bp). The following amplification protocols were used : (TTL) 95°C for 5 min, 35 cycles of [95°C for 1min / 50°C for 1 min / 72°C for 1 min], 72°C for 2 min; (SVBP) 95°C for 5 min, 33 cycles of [95°C for 30 sec / 50°C for 30 sec / 72°C for 30 sec], 72°C for 2 min. DNAs were analyzed on 1,2 % and 2 % agarose gels for TTL and SVBP, respectively.

### Immunoblotting

Cortical neurons were cultured for 15 days in vitro, lysed in Laemmli buffer and boiled at 96°C for 5 minutes. The total protein contents were equilibrated using stain-free 4%-15% gels (Bio-Rad) and then quickly transferred to Nitrocellulose using a Trans-Blot Turbo Transfer System (Bio-Rad). Immunoblots were developed using specific primary antibodies against detyrosinated α-tubulin (1:8000), tyrosinated α-tubulin (1:5000) and total α-tubulin (1:8000). Specific fluorescent secondary antibodies conjugated to Alexa 488 or Cy5 (Jackson Laboratories) were used and analyzed with a ChemiDoc™MP Imaging System (Bio-Rad) using Image Lab software (stain-free gel and fluorescence protocol) for quantification. For each lane of the blot, the software measures the integrated intensity of the band corresponding to the antigen of interest. The signal of each modified tubulin (tyrosinated or detyrosinated) was normalized to the signal of total α-tubulin of the same lane and the ratio of normalized signals was calculated. One neuronal culture per embryo was processed as indicated and for each neuronal culture, 3 independent blots were performed. For detailed information about antibodies, see Supplementary Tables.

### Wildtype mouse culture for TTL shRNA knockdown

P0 “wildtype” C567BL/6J mouse pups were decapitated, the brains were dissected in ice-cold HBSS (Sigma, H9269), and the hippocampi of several animals were collected in 450 µl cold HBSS. 50 µl trypsin (Gibco, 12499-015) were added to a final concentration of 0.025% and incubated for 15 min at 37°C. After 2 x washes with 500 µl warm HBSS, the hippocampi were resuspended in 1 ml warm plating medium (DMEM (Gibco, 41966-029) with 10% FBS and 1% Penicillin/Streptomycin (Invitrogen, 15140122)) and mechanically dissociated by carefully pipetting up and down through two different needles with different pore sizes (yellow 20G and brown 26G) with a 2 ml syringe, up to 4 times each. The cells were counted manually using a Neubauer chamber by staining 15 µl of the cell suspension with trypan blue. For microfluidic chambers, 70.000-80.000 cells were centrifuged at 1000 g for 2 min, the supernatant was removed, and the cells were resuspended in 5 µl plating medium, and injected into the microfluidic chamber. The four wells of the chamber were then filled up with plating medium, and the cultures were incubated at 37°C with 5% CO2 for 2 hours before the culture medium was replaced with growth medium (Neurobasal A (Thermo Fisher, 12349015) supplemented with 1 x B27 Plus (Thermo Fisher, A3582801), 4 mM Glutamax (Gibco, 35050061) and 1 mM sodium pyruvate (Gibco, 11360070)). The cells were then transduced on DIV5 by the addition of AAV, containing either TTL shRNA or control shRNA expression vectors, directly into the cell medium (see Supplementary Tables). 7-8 days after transduction (DIV12-13), neurons were imaged with Lysotracker Red as described below.

### Dual imaging of Magic Red and Lysotracker Green

Magic Red™ was reconstituted with DMSO, aliquoted and stored at −80 °C according to the manufacturer’s instructions. Primary hippocampal neurons were prepped from C57BL/6 P0 mice. 80 000 cells were plated into the somatodendritic compartment of a 2-compartment microfluidic chamber (MFC). At DIV14 the MFC was placed in a stage top incubator (okolab) at 37 °C, 5 % CO2 and 90 % humidity atmosphere at the microscope. LysoTracker™ Green DND-26 (ThermoFisher Scientific, Invitrogen™, L7526) and Magic Red™ (Biomol, Immunochemistry Technologies, ICT-937) were diluted in the conditioned medium to final dilutions of 1:10.000 for LysoTracker™ and 1:250 for Magic Red™ and added back to both the somatodendritic and axonal compartment of the MFC. After short incubation, confocal imaging was performed with a Nikon Eclipse Ti-E Visitron SpinningDisk confocal microscope controlled by VisiView software (Visitron Systems). The samples were imaged using a 100× TIRF objective (Nikon, ApoTIRF 100×/1.49 oil) resulting in a pixel size of 65 nm with 488 and 561 nm excitation laser. Lasers were coupled to a CSU-X1 spinning-disk (Yokogawa) unit via a single-mode fiber. Emission light was collected through a filter wheel with filters for GFP (Chroma ET525/50m) and RFP (ET609/54m). Z-stacks were acquired with a pco.edge 4.2 bi sCMOS camera (Excelitas PCO GmbH) with 350 nm step size in 16-bit depth. For image analysis, the FIJI plugin “ComDet” (v.0.5.5) was used to detect particles in the green channel (Lysotracker-positive organelles). The detected ROIs were then used to measure fluorescence intensity in both the green and red channels.

### Immunocytochemistry

Cells were fixed in 4% Roti-Histofix (Carl Roth), 4% sucrose in PBS for 10 min at RT, and washed three times with PBS, before they were permeabilized in 0.2% Triton X-100 in PBS for 10 min. The cells were then washed 3× in PBS and blocked for 45 min at RT with blocking buffer (BB/ 10% horse serum, 0.1% Triton X-100 in PBS). Incubation with primary antibodies was performed in BB at 4°C overnight. After 3× washing in PBS, cells were incubated with corresponding secondary antibodies in BB for 1 h at RT and washed again 3× 10 min in PBS. If phalloidin-staining was included, the coverslips were subjected to a second overnight incubation step with 1:100 phalloidin-Atto647N (Sigma, 65906) in PBS and washed again 3× 10 min in PBS. As a final step, coverslips were post-fixed in 2% Roti-Histofix, 2% sucrose in PBS for 10 min at RT, washed 3× 10 min with PBS, and mounted on microscope slides using Mowiol. Mowiol was prepared according to the manufacturer’s instructions (9.6 g mowiol 4–88 (Carl-Roth, 0713.1), 24.0 g glycerine, 24 ml H_2_O, 48 ml 0.2 M Tris pH 8.5, 2.5 g DABCO (Sigma-Aldrich, D27802)).

### Imaging of AIS markers

Z-stack images of fixed neurons were acquired on Leica TCS SP8 and Leica TCS SP5 confocal microscopes with 488 nm, 568 nm and 633 nm excitation lasers using a 63.0×1.40 NA oil objective. The pixel size was set to 90 nm and Z-steps varied between 250 and 350 nm.

### Live imaging of Lysotracker in knockout and wildtype neurons

Live imaging of knockout cultures grown in MFC was conducted on DIV12. Lysotracker Red DND-99 (ThermoFisher Scientific, L7528) was diluted in the conditioned medium to a final dilution of 1:10.000 and added back to the somatodendritic compartment of the MFC. Lysotracker-stained vesicles became visible in the axonal compartment a few minutes later. Imaging was conducted at 37°C and 5% CO2 on a Zeiss Axio Observer coupled to a spinning disk confocal system (CSU-W1-T3; Yokogawa) connected to an electron-multiplying CCD camera (ProEM+1024, Princeton Instruments). Images were taken every 1 sec for 180 sec with a 63× oil immersion objective (1.46 NA). STED imaging was done on an Abberior 2D-STEDYCON module installed on a conventional epifluorescence microscope (Zeiss) with a 100× oil immersion objective (1.46 NA).

### Imaging of Lysotracker in TTL knockdown neurons

Live imaging of TTL shRNA knockdown cultures grown in MFC was conducted on DIV12-13. Lysotracker Red DND-99 (ThermoFisher Scientific, L7528) was diluted in the conditioned medium to a final dilution of 1:10.000 and added back to the somatodendritic compartment of the MFC. Lysotracker-stained vesicles became visible in the axonal compartment a few minutes later. shRNA-transfected axons were identified via GFP expression. Imaging was conducted at 37°C and 5% CO2 with a spinning disc confocal microscope (Nikon ECLIPSE Ti) controlled by VisiView software (Visitron Systems) and equipped with the following components: Spinning Disk (Yokogawa), solid-state lasers (488, 561, 647, 405), an EM-CCD camera (Hamamatsu, Digital Camera C9100), and with a 100 × objective (NA 1.45). The image acquisition rate was 1 fps over 3 min.

### Semi-automated analysis of kymographs

Axons of Lysotracker-treated neurons grown in MFC were imaged as above. Timelapse image stacks were processed using FIJI/ImageJ. For shRNA-transfected neurons, individual axons were identified by their GFP expression. For knockout neurons, individual axons were identified by a brightfield image taken before live imaging. Only axons where the directionality could be established (i.e. axons directly coming out of the MFC’s microgrooves) were chosen. Individual axons were manually traced using the segmented line tool with a 10-pixel width (pixel size = 0.175 µm). Kymographs were then generated from those ROIs using the KymoResliceWide plugin (https://github.com/UU-cellbiology/KymoResliceWide). The kymographs were then used as input for the bidirectional KymoButler deep learning software (Jakobs et al., 2019, https://github.com/elifesciences-publications/KymoButler) run in Wolfram Mathematica with the following settings: detection threshold=0.2, decision threshold=0.5, minimum particle size=30, minimum frame number=20. Automatically detected traces from KymoButler were loaded into a self-made MATLAB program to overlay them with the original kymograph image, and manually corrected when there was a visible error. The researcher conducting the manual correction was blinded to the data’s identities. The corrected tracks were then exported as lists of x-y coordinates, and a self-made Python-based program was used to extract various parameters from the data (see Fig. S2; https://github.com/HU-Berlin-Optobiology/AIS-project.git). In short, the program assigned “movement” if the x-coordinate (x-axis = space) changed from one frame to the next (y-axis = time), and assigned “no movement” if the x-coordinate stayed the same from one frame to the next. Cutoff values were then set to identify “runs” as a continuous movement for 5 frames, and “pause” for continuous non-movement for 5 frames (i.e. a “pause” was defined as a period in which a particle defined as mobile, which had at least one processive run during the imaging period, did not move for 5 or more consecutive frames). Anything below those cutoffs was assigned as “oscillating”. A track that did not contain any runs was identified as completely stationary. “Runs” were further divided into anterograde runs and retrograde runs. For each individual identified track, the program calculated the duration of runs and pauses, the run distance and run velocity, and exported those values both individually and as averaged values per each track. Only the run distance and velocity of periods where the particles underwent processive runs for at least 5 consecutive frames were taken into account. For each mobile particle, the net displacement in anterograde or retrograde direction was determined. If a particle moved both in the retrograde and anterograde direction during the imaging period, it might end up having no net displacement (“none”) despite being mobile. The threshold for net displacement was set to 5 pixels (0.875 µm).

### Statistical analysis

All statistical analysis was conducted in GraphPad Prism 7.05. Data are shown as mean values with standard error of the mean (SEM). Data was tested for Gaussian distribution using the D’Agostino & Pearson normality test and accordingly subjected to parametric or non-parametric statistical tests. Statistical analysis of differences between two groups was performed using Student’s t-tests for populations with Gaussian distribution, or Mann-Whitney’s test for non-Gaussian distributions. When comparing 3 or more univariate samples we used one-way ANOVA, or the non-parametric Kruskal-Wallis test. Post-hoc comparisons following Kruskal-Wallis test were done with the non-parametric Dunn test.

## Acknowledgements

We would like to thank Eitan Zahavi (Weizman Institute of Science, Rehovot) for providing molds for microfluidic devices, Alexander Biermeier (AG Optobiology, HU, Berlin) for help with their production, Julia Bär for help with prepping primary neurons, Matthias Kneussel (ZMNH, Hamburg) for access to the spinning disc confocal system, Frederic Saudou and Anca Radu (GIN, Grenoble) for help with making MFCs and for access to equipment, and Virgilio Failla (UMIF, UKE, Hamburg) for access to the laser scanning confocal microscope. This work was supported by the Photonic Imaging Center of Grenoble Institute Neuroscience (PIC-GIN, Univ Grenoble Alpes – Inserm U1216) which is part of the ISdV core facility and certified by the IBiSA label. We would like to thank Isabelle Jacquier (GIN, Grenoble) for technical help in the preparation of primary neuronal cultures and genotyping, and Yasmina Saoudi from PIC-GIN, Grenoble, for helping us with confocal and STED microscopy.

## Data availability

N/A

